# A High-Throughput Gravimetric Phenotyping Platform for Real-Time Physiological Screening of Plant–Environment Dynamic Responses

**DOI:** 10.1101/2020.01.30.927517

**Authors:** Ahan Dalal, Itamar Shenhar, Ronny Bourstein, Amir Mayo, Yael Grunwald, Nir Averbuch, Ziv Attia, Rony Wallach, Menachem Moshelion

## Abstract

Food security for the growing global population is a major concern. The data provided by genomic tools far exceeds the supply of phenotypic data, creating a knowledge gap. To meet the challenge of improving crops to feed the growing global population, this gap must be bridged.

Physiological traits are considered key functional traits in the context of responsiveness or sensitivity to environmental conditions. Many recently introduced high-throughput phenotyping techniques are based on remote sensing or imaging and are capable of directly measuring morphological traits, but measure physiological parameters only indirectly.

This paper describes a method for direct physiological phenotyping that has several advantages for the functional phenotyping of plant–environment interactions. It aims to help users overcome the many challenges encountered in the use of load-cell gravimetric systems and pot experiments. The suggested techniques will enable users to distinguish between soil weight, plant weight and soil water content, providing a method for continuous and simultaneous measurement of dynamic soil, plant and atmosphere conditions, alongside key physiological traits. This method allows researchers to closely mimic field stress scenarios while taking into consideration the environment’s effect on the plant’s physiology. This method also minimizes pot effects, which are one of the major problems in pre-field phenotyping. It includes a feedback fertigation system that enables a truly randomized experimental design with a field-like plant density. This system detects the soil-water content limiting threshold (θ) and allows for the translation of data into knowledge through the use of a real-time analytic tool and an online statistical resource. This method for the rapid and direct measurement of the physiological responses of multiple plants to a dynamic environment has great potential for use in screening for beneficial traits associated with responses to abiotic stress, in the context of pre-field breeding and crop improvement.

**SUMMARY:** This high-throughput, whole-plant water relations gravimetric phenotyping method enables direct and simultaneous real-time measurements and analysis of multiple yield-related physiological traits involved in dynamic plant–environment interactions.

## INTRODUCTION

Ensuring food security for a growing global population under deteriorating environmental conditions is one of the major goals of agriculture research (Ray et al., 2013; FAO, 2017; Dhankher and Foyer, 2018). The availability of new molecular tools has greatly enhanced crop-improvement programs. However, while genomic tools provide a massive amount of data, our limited understanding of the actual phenotypic traits creates a great knowledge gap. Bridging this gap is one of the greatest challenges facing modern plant science (Chen et al., 2014; Ubbens and Stavness, 2017; Danzi et al., 2019). To meet the challenges that arise in crop improvement and minimize the genotype–phenotype knowledge gap, we must balance the genotypic approach with a phenocentric one (Miflin, 2000; Dalal et al., 2019).

Recently, various high-throughput phenotyping (HTP) platforms have allowed for nondestructive phenotyping of large plant populations over time and these platforms may help us to reduce the genotype–phenotype knowledge gap (Moshelion and Altman, 2015; Singh et al., 2016; Dalal et al., 2019; Danzi et al., 2019). Most of these HTP platforms involve the assessment of phenotypic traits through electronic sensors, robotics and/or automated image acquisition (Tardieu et al., 2017; Negin and Moshelion, 2017). However, the reliability and accuracy of currently available imaging techniques for in-depth phenotyping of dynamic genotype–environment interactions and plant stress responses are questionable (Li et al., 2014; Gosa et al., 2018).

In this paper, we demonstrate a nondestructive method involving an HTP platform that includes weighing lysimeters (a gravimetric system) and environmental sensors. This system can be used for the collection and calculation of a wide range of data, such as whole-plant biomass gain, transpiration rates, stomatal conductance, root fluxes and water-use efficiency (WUE). Rice (*Oryza sativa* L.) was used as a model crop and drought was the examined treatment. Rice was chosen as it is a major cereal crop with wide genetic diversity and it is the staple food for over half of the world’s population (reviewed by Ito and Lacerda, 2019). Drought is a major environmental abiotic stress factor that can impair plant growth and development, leading to reduced crop yields (reviewed by Anjum et al., 2011). This crop-treatment combination was used to demonstrate the platform’s capabilities and the amount and quality of data that it can produce. For more information regarding the theoretical background for this method, please see Halperin et al. (2017).

## PROTOCOL

### 1. Get to know the system components

#### 1.1. Weighing lysimeters

The lysimeter includes the load cell, which converts the mechanical load of an object into an electrical charge, and a metal platform that covers the upper and lower parts of the load cell, so that the object’s weight can be properly measured. The lysimeter is covered with a polystyrene block and a plastic cover for heat insulation (**Figure 1A,B**).

**Figure 1.**
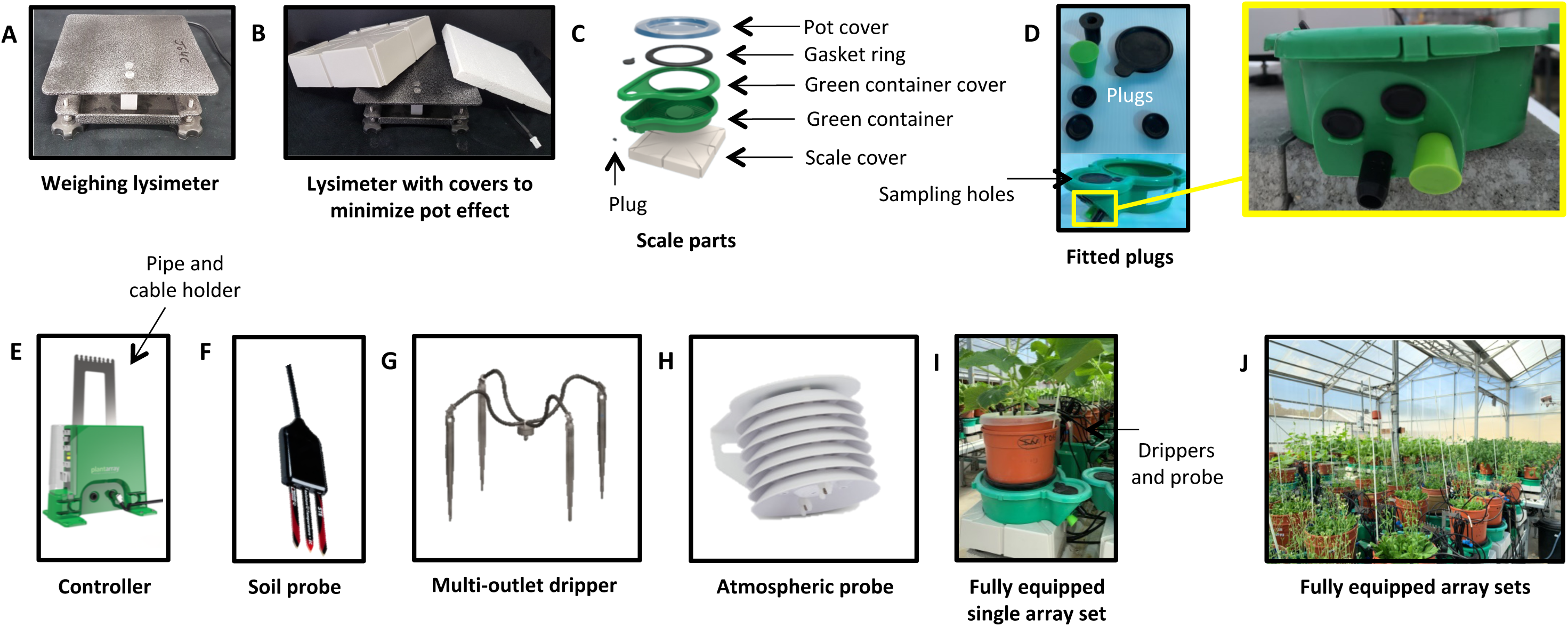
Components and set up of the gravimetric phenotyping system.

#### 1.2. Planter unit container

A water reservoir (green container) is placed on the lysimeter cover to collect the liquid that drains from the pot. The container has a drainage extension with four holes (with plugs) at different heights, which can be used to adjust the water level in the container after the drainage through a particular hole stops (the reserve water volume). The desired water volume will depend on the plant species, the type of growing media and the water requirements of the plants (i.e., estimated daily transpiration volume).

The green container is coupled with a green cover, which has a large round opening through which the pot is inserted. A black rubber gasket ring is attached to one side of the green cover and the pot is attached to the other side, to minimize the amount of water lost to evaporation from the container. The green cover has two sampling holes (small and big) above the drainage extension, which are sealed with rubber plugs (**Figure 1C,D**).

#### 1.3. Control unit

The control unit consists of a green rectangular box that contains the electronic controller and solenoid valves. There are holes through which fertigation solution can enter and exit the pots, as well as sockets for connecting the load cell and different sensors. Different treatments, such as different levels of salinity or different mineral compositions, can be applied via the fertigation solution. A metal stand is connected to the controller, to hold the pipes and cables and prevent them from touching the pots and adding weight **(Figure 1E)**.

#### 1.4. Probes and drippers

The other components required are soil probes (e.g., moisture, temperature and EC sensors - 5TE; **Figure 1F**), multi-outlet drippers (for fertigation and/or treatment applications; **Figure 1G**) and atmospheric probes [for measuring vapor pressure deficit (VPD) and radiation; **Figure 1H**]. For an overview of a fully equipped single array set-up, please see **Figure 1I,J**.

#### 1.5. Parts required for a single pot set-up

The following components are needed: one 4-L pot, one 4-L pot that does not have bottom (to serve as a net holder), one circular piece of nylon mesh (pore size = 60 mesh) with a diameter double that of the bottom of the pot, one cover with designated holes for plant and irrigation drippers, one 60-cm white fiberglass stick (pole) and one black gasket ring (**Figure 2A,B**).

**Figure 2.**
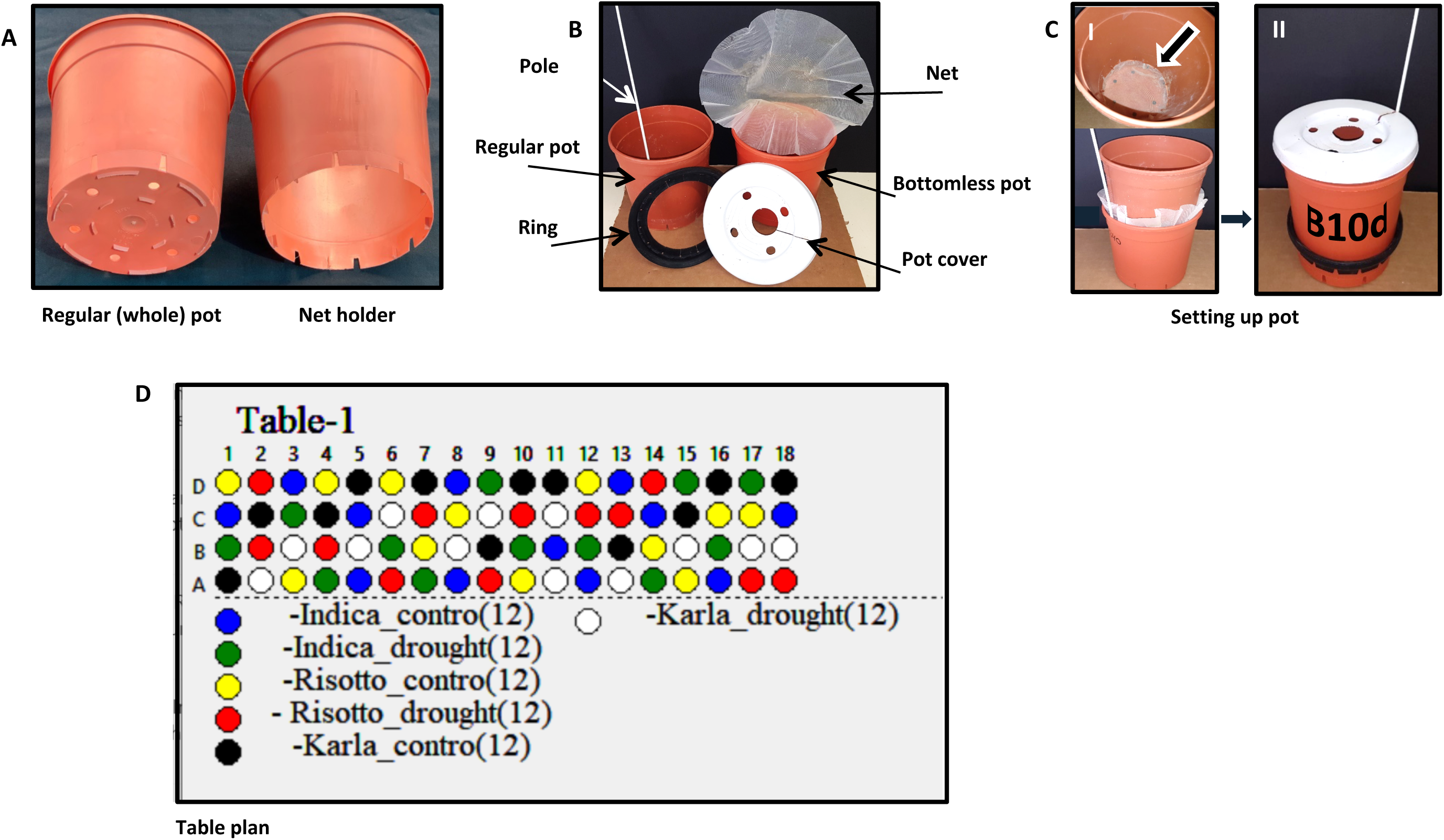
(A-C) Components for pot preparation. (D) Example for a randomized pots table plan.

### 2. Prepare pots for an experiment

2.1. (A). Spread the nylon mesh on top of the whole pot and place the net holder on top of the mesh. Slowly push the net holder half way down the inside of the whole pot. Make sure that the mesh remains uniformly spread as it is pushed down between the two pots. (B). Insert the fiberglass stick between the two pots and push it all the way down to the bottom of the whole pot, making sure it is on the outer side of the mesh as well. Before pushing the net holder all the way down, (C) push the mesh down by hand from inside the pot and adjust so that it is spread uniformly over the bottom of the pot once the net holder has been fully inserted (**Figure 2CI**).
2.2. Slide the gasket ring from the bottom of the pot set-up described above, a third of the way up the side of the pot. Make sure that the slits of the ring open toward the bottom of the pot (**Figure 2CII**).
2.3. Repeat steps 2.1 and 2.2 for all of the experimental pots before continuing on to the next step. Randomize the location of your plants (either randomized block design, or fully random design) using the Array Randomizer application. (To download the free program and for more information, please see the link in the supplementary Excel file.) Label the pots according to their physical position in the array in the greenhouse. For example, the label “B10D” corresponds to a pot located on Table B in Column 10 and Row D. In our greenhouse, each table has 1–18 columns and four rows (**Figure 2D**), yet the array structure can be easily adjusted to any shape based on the size of your own greenhouse. Prepare three additional pots for each table for soil water content measurements (please see 8.1).

### 3. Grow your plants

3.1. Choose your growing medium smartly. Selecting the most suitable growing medium for the gravimetric system experiment is one of the crucial steps. In principle, it is recommended to choose the soil media which have fast drainage properties, rapid arrival to pot capacity and high pot capacity stability, as these features allow more accurate measurements by the gravimetric system. In addition, different treatments designed in the experiment must also be considered. For example, treatments with salts, fertilizers or chemicals require an inert substrate, preferably with low cation exchange capacity. Fast drought treatment for a low transpiring plant will benefit from soils with low VWC, and vice versa. If the roots are required for a post-experiment analysis (e.g. root morphology, dry weight, etc.) then media with lower organic matter content (i.e. sand, porous ceramic or perlite) will be easier to wash with less damage to the root system. For an experiment planned for longer period, it is recommended not to choose media that are rich in organic matter as it may get degraded with time. Please see **Table 1** and **2** for more details, which may serve as a guideline to choose the right medium for the experiment.

**Table 1.**
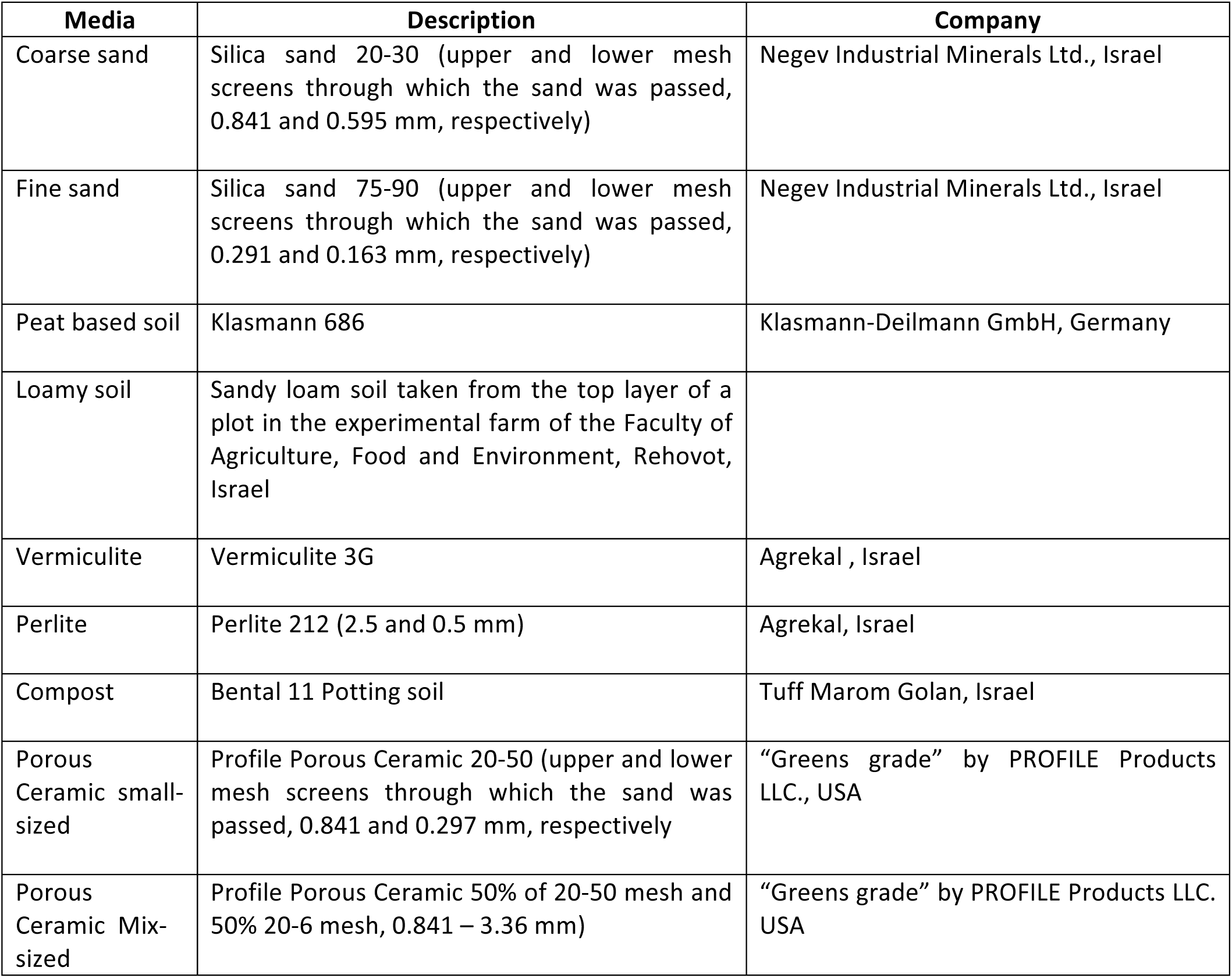
Media components.
3.2. Germinate your seeds in the desired potting medium/soil in trays if possible, on side tables inside the greenhouse, in order to acclimatize the plants to the environmental conditions in the greenhouse.
3.3. If seedlings are not initially germinated in trays, after germination, transfer the seedlings into cavity trays with potting medium/soil, one seedling per cavity, and grow them until their roots are dense enough to take the shape of the cavity (root-soil plug).
3.4. Leave 5–7 cavities without seedlings for soil weight measurements (only potting medium/soil; **Figure 3**). For more information, please see section 6.7.

| **Media** | **Description** | **Company** |
| --- | --- | --- |
| Coarse sand | Silica sand 20-30 (upper and lower mesh screens through which the sand was passed, 0.841 and 0.595 mm, respectively) | Negev Industrial Minerals Ltd., Israel |
| Fine sand | Silica sand 75-90 (upper and lower mesh screens through which the sand was passed, 0.291 and 0.163 mm, respectively) | Negev Industrial Minerals Ltd., Israel |
| Peat based soil | Klasmann 686 | Klasmann-Deilmann GmbH, Germany |
| Loamy soil | Sandy loam soil taken from the top layer of a plot in the experimental farm of the Faculty of Agriculture, Food and Environment, Rehovot, Israel |  |
| Vermiculite | Vermiculite 3G | Agrekal , Israel |
| Perlite | Perlite 212 (2.5 and 0.5 mm) | Agrekal, Israel |
| Compost | Bental 11 Potting soil | Tuff Marom Golan, Israel |
| Porous Ceramic small-sized | Profile Porous Ceramic 20-50 (upper and lower mesh screens through which the sand was passed, 0.841 and 0.297 mm, respectively) | “Greens grade” by PROFILE Products LLC., USA |
| Porous Ceramic Mix-sized | Profile Porous Ceramic 50% of 20-50 mesh and 50% 20-6 mesh, 0.841 – 3.36 mm) | “Greens grade” by PROFILE Products LLC. USA |

**Figure 3.**
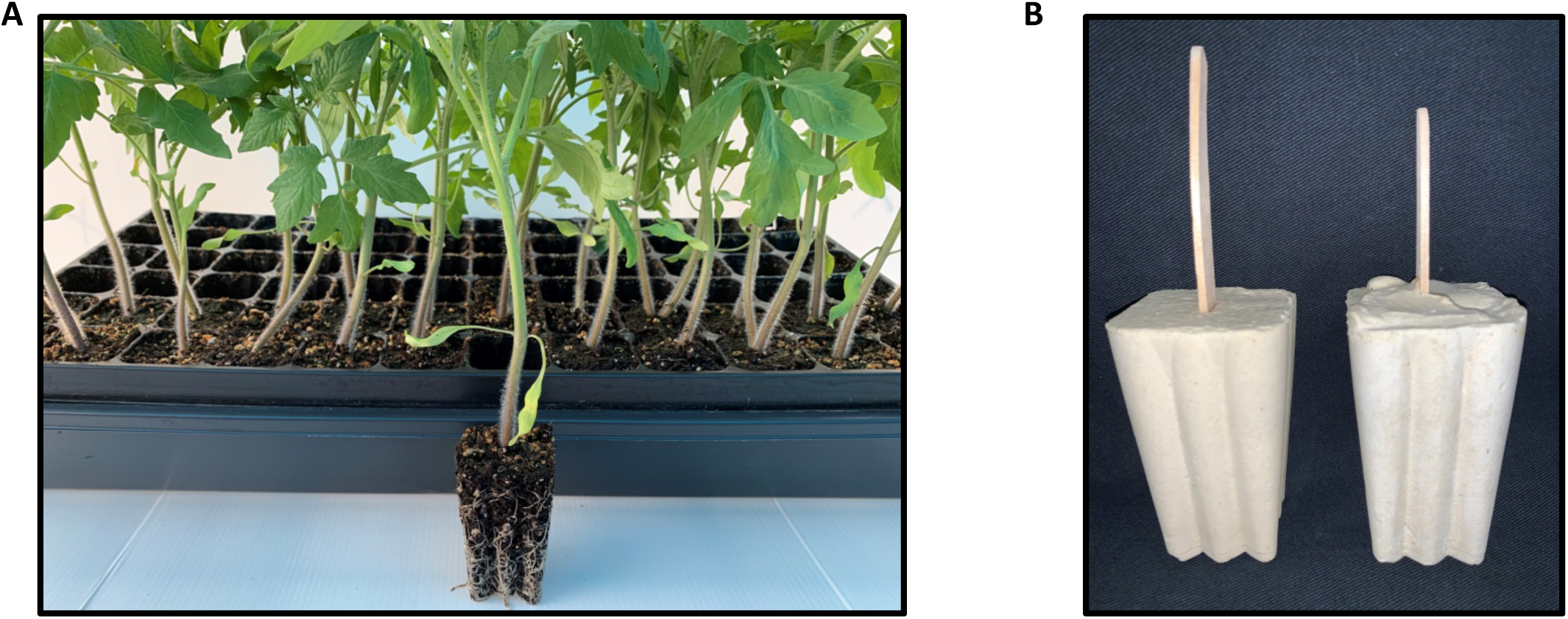
A) Plants growing in cavity trays. B) Casts of cavity-mold for creating cavities in the soil/media for easy transplanting

### 4. Calibrate the lysimeter

Check that all of the lysimeters are level using a spirit level and then start the weight calibration process. Use two standard weights between 1–10 kg. Perform the calibration while the green container, including all plugs, is on the load cell.

Follow the steps below:

4.1. Put the first (lighter) calibration weight on each load cell.
4.2. In the operating software, go to the Calibration tab and choose the weight for your first point. Then, select the load-cell position where you placed the weight and click “Get point1” (**Figure 4A**).

**Figure 4.**
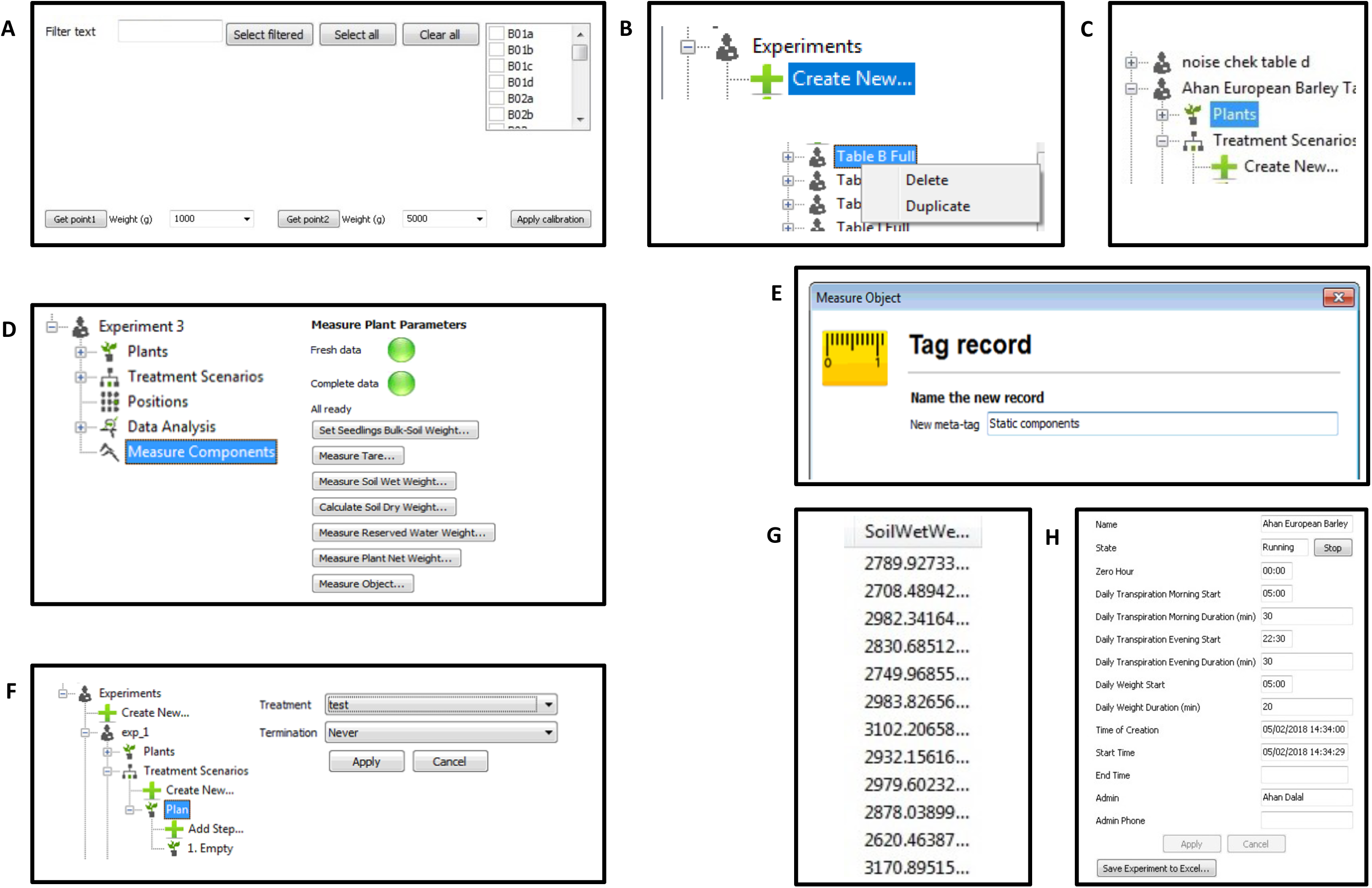
Operating software windows for setting up experiment (Plantarray 3.0 system; Plant-DiTech)
4.3. Repeat for the second weight and click “Get point2”.
4.4. Click “Apply calibration”.

### 5. Set up the experiment

The process of setting up the experiment is designed to take into account the weight of all the parts of the system, namely, the weight of the growth medium weight (including the soil water weight at pot capacity) and the initial weight of the seedlings. Weight measurements are taken separately for tare (including pot sets, soil probes and all plastic parts), wet soil (pot capacity), dry soil, reserve water in the containers, and seedlings.

Follow the steps below.

5.1. To start a new experiment, open the operating software. Open the Experiments tab in the menu on the left side of the screen. Click on Create New or duplicate the experiment properties from a previous experiment by right-clicking on the desired experiment and choosing “Duplicate”. Rename your experiment as: “Experimenter name, Experiment name, table or start date” (**Figure 4B**).
5.2. Make sure that no unit is being used in a different experiment. Check that all plants in the Plants table match your experimental design. If not, change the table according to your design (please see section 7; **Figure 4C**).
5.3. Start the experiment by clicking on the experiment name and then click “Start”.
5.4. Take manual measurements of the pre-prepared empty pots (double pot, net, stick and black gasket ring).
5.5. Mix the potting medium/soil thoroughly with sufficient water so that it breaks down into homogeneous particles and is saturated, to ensure uniformity and homogeneity. * We recommend using a mechanical mixer (e.g., a concrete mixer). * When a highly homogenous medium (i.e., industrial sand) is being used, this step can be skipped.
5.6. Fill all of the pots for the experiment with the desired potting media (e.g., sand/soil/peat).
5.7. Insert a cast of a cavity mold (**Figure 3B**) of a similar shape and size as that of the root-soil plug of the seedlings (from cavity tray) in the middle of the potting medium and push it in completely. Tap on the bottom of the pot a few times to make sure the potting medium is well distributed in the pot. Repeat for all pots. Next, irrigate the pots well and rinse off the outside of the pots. Allow the pots to drain for 30 min before continuing on to the next step. Make sure that the pots drain quickly. If the potting medium drains too slowly (e.g., dense peat), premix it with an airy substance (e.g., perlite, please see also **Table 1** and **2**). Removing the cast (step 6.4 below) will leave a cavity that accurately fits the root-soil plug of the seedlings, which will ensure the successful transplanting of the seedlings into the pots.
5.8. Place all of the filled pots on the lysimeter array (in the green containers that are already there) according to your experimental design (**Figure 2A**).
5.9. Check that the green containers are well-fitted into the load cell cover and not touching one another.
5.10. In the operating software, open your experiment tab and select the Measure Components tab. Click on “Measure object”. Name the measurement “1^st^ measurement” (**Figure 4D**).
5.11. Place the irrigation drippers, probes and pot covers on each pot. Make sure that the lines for the multi-outlet drippers and the probe cables are supported by their respective stands (attached to the units for each lysimeter scale) before placing them in the pots.
5.12. Wait for a few minutes for a new measurement to be taken (data are collected automatically every 3 min) and then open your experiment tab. Select the Measure Components tab and click “Measure Object”. Meta-tag this measurement to the “1 ^st^ measurement” you took before and name it “Static components” (**Figure 4E**). Meta-tags are used when you want to record a value that is determined by subtracting one measured value from another. NOTE: After making any adjustments to the system, wait for a new data point to be recorded (every 3 min) before taking the next measurement.
5.13. Check the Static Components column to confirm that the values recorded in the Plants table do not include outliers. If the numbers recorded are too low or too high, check for any interference with the load cell (e.g., that nothing is touching it), and then take a new measurement.
5.14. Extract the Plants table as an Excel file, add the average pot weight (from step 5.5) to the measurement of the static components and label the column “Tare weight”. Upload the file.
5.15. Make sure that all of the drippers are securely inserted into the potting medium. Back in the operating software, in your Experiment tab, select “Treatments Scenarios”. Click “Create New” to make a new “Plan” and open it. In the plan, choose the first step (create a new step if needed) and open it. Choose “Test” for Treatment and “Never” for Termination. In the step option, you can choose any treatment that is listed in the Irrigation Treatments tab above “Experiments” (**Figure 4F**, please see also step 5.17).
5.16. Extract the Plants table as an Excel file, add “Plan” to the Treatment column and add “1” to the Step column. Upload the file.
5.17. Under the Irrigation Treatments tab, choose the “Test” treatment and set it to an irrigation time of 250 s. (Set the time 2 min ahead and go to the table in the greenhouse). Other treatments can also be created (please see step 8.3).
5.18. Check visually that all of the drippers are working and that water is dripping out of the perforated drain plug of the green container.
5.19. In your experiment, change the irrigation treatment on Plan “X”, Step 1 (please see step 5.16), to “normal irrigation treatment”. Make sure that each night irrigation is divided into several short pulses (events) with substantial pause between them (at least threeevents every night), to ensure that the soil reaches its field capacity.
5.20. Let the irrigation program run for 1 or 2 days to let the soil reach its field capacity and continue on to the next phase.

### 6. Start the experiment

**Note that the data collected at this stage will be used as a reference point for the rest of the experiment. Therefore, it is important to follow the next steps carefully.**

6.1. Repeat steps 5.15. and 5.16. (Alternatively, you can start the process in the early morning not long after the latest irrigation step.)
6.2. Check visually that all the pots are irrigated and that excess irrigation liquid is dripping out of the perforated drain plug.
6.3. Remove the green unperforated plug (at the lowest orifice) of the green container and let the water drain out completely. Then, put the plug back in its place (**Figure 1D**).
6.4. In the operating software, open the tab for your experiment and go to “Measure Components”. Click “Measure Object” and name the measurement as “Cast-pre”. Gently remove all of the casts from the pots and then wait 3 min for a new measurement to be recorded (**Figure 4D**). Click “Measure Object”, name it “Cast-post” and meta-tag the measurement to “Cast-pre”. The program will automatically calculate the difference between the two measured values and give you the cast weight to verify the weight sensitivity.
6.5. Check the weight measurements in the Plants table. The difference between the “Cast-post” measurements should not be more than 20 or 30 g.
6.6. To measure the weight of the wet soil, in the operating software, go to the Measure Components tab in your experiment and select the Measure Soil Wet Weight option. Take the measurement by clicking “OK” when asked. Check the Soil Wet Weight measurements in the Plants table of your experiment. The weight will appear in the Soil Wet Weight column (**Figure 4D,G**). If some of the measurements seem to fluctuate inappropriately, please do the following:
  a. Confirm that the pots are placed correctly and are not touching any neighboring pot(s).
  b. Disconnect the first pot on the table (the first pot that is connected to the electricity) for 2 min and then reconnect it.
6.7. Measure the average (3–5 repetitions) weight of a few empty cavities (from 3.3) manually. Press “Set Seedling Bulk-Soil Weight” and fill in the average weight (**Figure 4D**).
6.8. Click “Measure Plant Net Weight”. This first measurement is a reference point before planting (**Figure 4D**).
6.9. Make sure that the seedlings in the cavity trays are well irrigated (to field capacity after drainage). Gently pull out the seedlings with the root-soil plug from the cavities, making sure not to injure them, and place them carefully into the cavities made by casts in the pots, according to the experiment design. It is preferable to transfer the plants at dawn or dusk.
6.10. Click “Measure Plant Net Weight” again. This second measurement is the plant net weight. Meta-tag the measurement to the first one (the reference point). The software will calculate the difference between the two measurements and subtract the Seedling Bulk-Soil Weight. The result is the plant net weight.
6.11. Check the measured values in the Plants table of your experiment to make sure that they fall in a reasonable and logical range (**Figure 4C**).
6.12. Saturate the soil by repeating steps 5.15 and 5.16.
6.13. Make sure that all of the pots are draining properly. If not, repeat saturation again. Wait 30 min for the drainage to cease.
6.14. Under the Measure Components tab, click Measure Reserved Water Weight (**Figure 4D**).
6.15. Extract the Plants table as an Excel file, subtract the measured Plant Net Weight and Seedling Bulk-Soil Weight from the reserved water weight measurement, and label the column “Reserve Water Inventory”. Upload the file (**Figure 4C**).
6.16. Confirm that the time period during which daily transpiration will be recorded is appropriate for the goals of your experiment. Fill in the values in the experiment general tab as appropriate for your project (**Figure 4H**). Zero hour: The time at which the software will check whether it needs to move to the next step in the treatment scenario. Daily transpiration values: Daily transpiration is calculated as the difference between two weight windows during the day, for all days. The daily transpiration start time is the time at which the software will begin to measure the average weight.
6.17. It is recommended to monitor the plants for 1–2 days before starting a new experiment (duplicate and rename the experiment).

### 7. Change the Plants table

Extract the Plants table as an Excel file and change the table according to your needs. **Do not change the Plant IDs, Names or Positions.** Upload the file.

Labeling (grouping) columns: To present or analyze (please see step 9) grouped plants based on common labels (e.g., treatment, line), add a new column and label starting with # (e.g., #Treatment). In this column, make a notation for each plant (e.g., for “#Treatment” label, mark the plants as drought, control, etc.; **Figure 5**).

**Figure 5.**
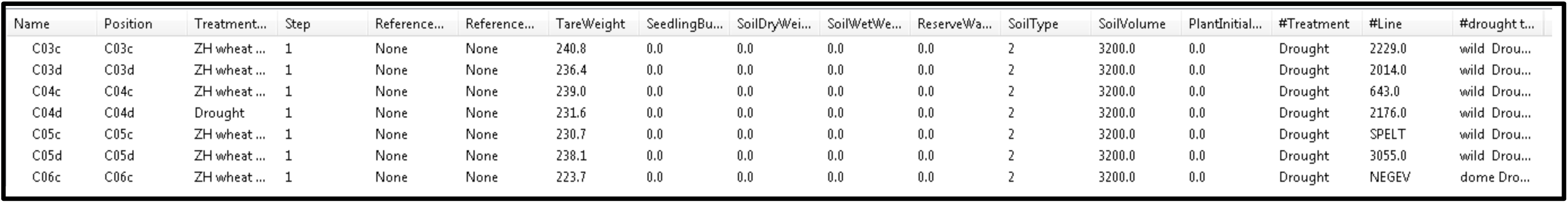
“Plants” table as MS excel file (Operating software)

### 8. Run the experiment

**8.1.** Calculating the soil gravimetrical water content / soil water content (SWC value).

NOTE: Gravimetric soil water content is different from volumetric soil water content (VWC). The SWC value is the ratio between the dry weight of the soil and the wet weight of the soil. To calculate SWC, use the three extra soil-filled pots without plants that were previously prepared and placed on a side table inside the greenhouse for a few days with regular irrigation. Weigh the wet soil in an aluminum tray in the early morning (as soon as possible after the last irrigation event).

Dry the aluminum tray with the soil in an oven (at 105 °C) for 4–5 days. Verify that the soil is completely dry by taking two consecutive weight measurements at least 30 min apart. If the weights are identical, the soil is indeed dry and the last measurement can be recorded as the dry soil weight.

In the operating software, go to Measure Components and click on the Calculate Soil Dry Weight tab. Fill in the soil wet and dry weights for each sample (**Figure 6**).

**Figure 6.**
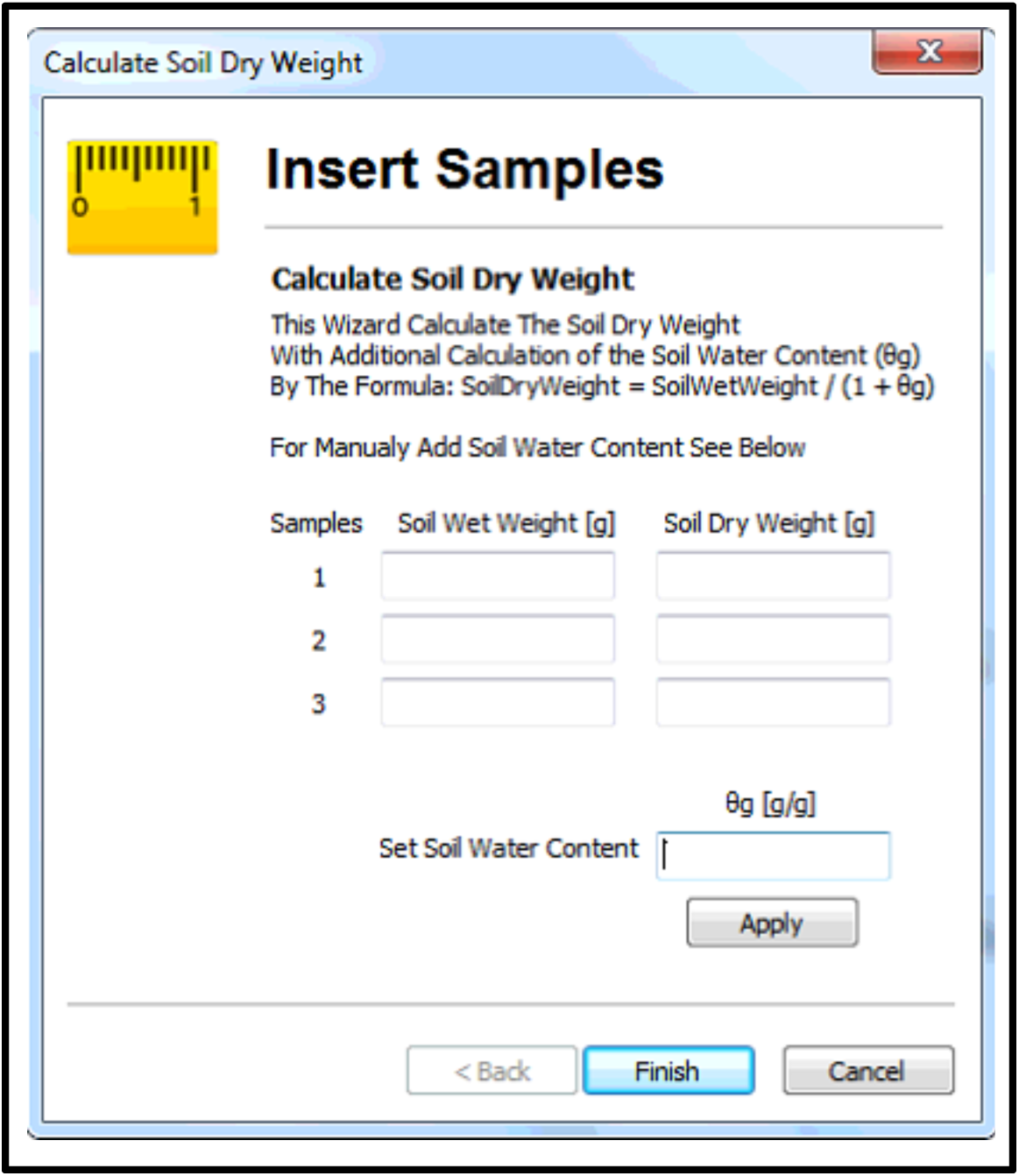
Operating software window for calculating “Soil Dry Weight”.

#### 8.2. Manual calculation of SWC

The equation shown below can be used to calculate the SWC value.

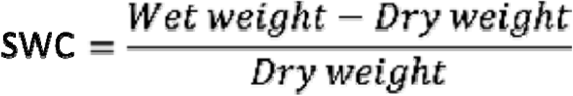

Average two measurements (SWC) from two pots. Select the Measure Components tab, enter the θg [g/g] value and click on “Calculate Soil Dry Weight. The soil dry weights of all of the experiment pots will be calculated automatically by the software (assuming that all of the pots in the experiment contain the same medium; **Figure 4D** and **Figure 6**).

#### 8.3. Application of irrigation treatments

Irrigation scenarios can be applied by composing a step-by-step treatment plan.

To compose a new irrigation treatment plan, go to “Irrigation Treatment”, click on “Create New” and name the new treatment. Open the specific treatment in the list of irrigation treatments and click the on the default “00:00”. In the main window (please see **Figure 7A**), “Time” indicates the time the valve will open (i.e., the beginning of the irrigation treatment). “Valve” is the valve to be opened (A or B, depending on the valve that is connected to the desired solution) and “Command Type” indicates the type of data that will be used to determine when the valve will be closed. “Command type” has four options:

**Figure 7.**
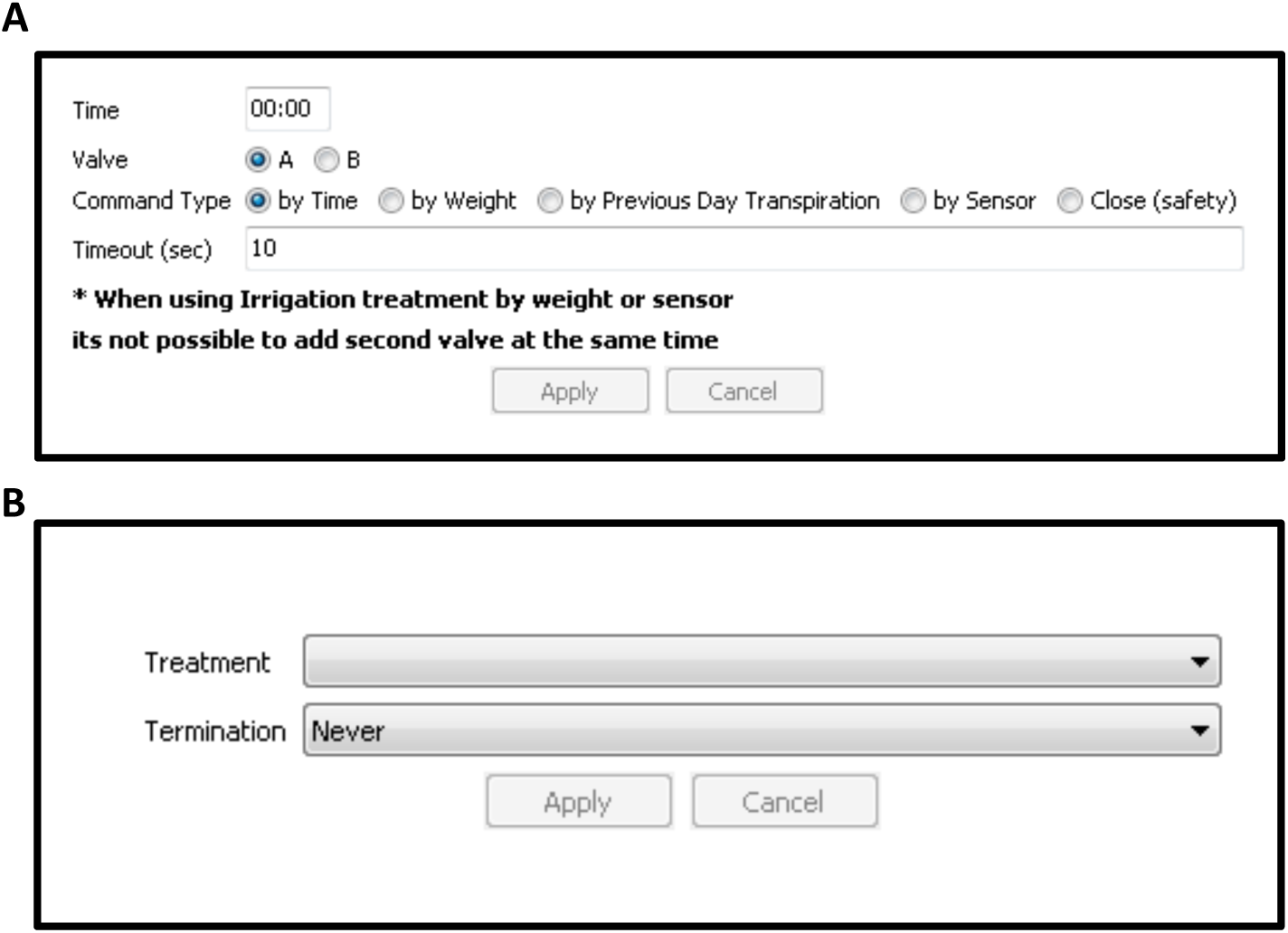
Operating software window for setting up an irrigation treatment.

1. By Time – How many seconds the valve will be open
2. By Weight – The weight gain/water (in grams) to be added to the pot via irrigation.
3. By Transpiration – Irrigation can be applied differentially to each pot based on the transpiration of each individual plant over the previous day. The user can decide what percentage of the previous day transpiration will be applied during irrigation. (Under the well-irrigated condition, it is suggested to give the plant more than 100%, in order to wash the soil and compensate for plant growth. Drought-treated plants should be given less water, with exact volumes based on the desired level of drought stress.)
4. By Sensors – Irrigation can be applied according to a sensor reading, such as apparent dielectric permittivity (which can be used to determine the VWC). Choose the sensor type, the desired parameter and the desired parameter value.

All possibilities include a Time Out option that will close the tap even if the set conditions were not reached. It is recommended to set the Time Out for a period longer than the set conditions. After defining the irrigation treatments for the experiment, open the desired experiment in the list of experiments, open “Treatment Scenario”, open default “Plan” and choose the first step (please see **Figure 7B**). In “Treatment”, choose an irrigation treatment from the list. Then, in “Termination”, choose the appropriate condition to stop the current step and move to the next one.

After selecting an irrigation scenario, open the experiment’s Plants table (**Figure 5**) and input the “Treatment” and “Step” for each plant. “Treatment” is the name of the treatment scenario and “Step” is the event number within the treatment scenario.

#### 8.4. Planning a drought treatment

Each individual plant has a unique transpiration rate based on its size and location in the greenhouse. To enable a standard drought treatment (i.e., similar drying rate for all pots during the treatment), it is recommended to plan a drought scenario and control it via the system’s feedback-irrigation tool. This feature enables the user to design irrigation for each individual pot, based on time, pot weight, data from a soil sensor (e.g., VWC) or plant transpiration over the previous day. Each plant can be irrigated individually in a customized manner based on its own performance. This differential irrigation minimizes the differences between the plants’ soil water contents, so that all of the plants are exposed to a controlled drought treatment regardless of their individual water demands (**Figure 8**).

**Figure 8.**
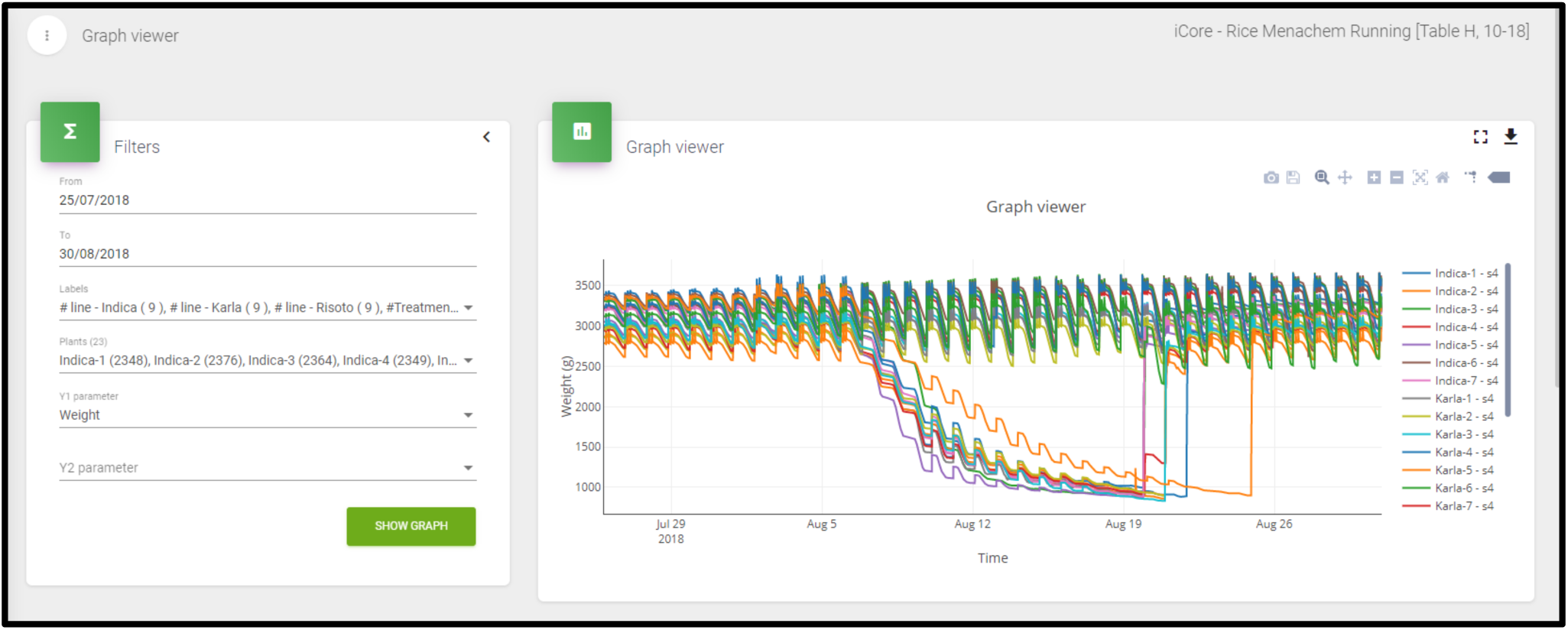
**“**SPAC Analytics” Graph viewer window. We used three varieties of rice, namely *Indica, Karla* and *Risotto*, treated with two different irrigation scenarios (i) well irrigated (control) and (ii) drought. The raw data represents variation in the weight of the plants during the experiment. Each line represents one plant/pot. During the day, the plant transpires, thus the system loses weight which is seen as slope in the line curves every day. The pots were irrigated every night (to pot capacity), represented as peaks in the line curves. The irrigation was followed by drainage to remove excess water after saturation. Initially, all plants were well irrigated (control); from Aug 07’ 2018, half of the plants were given drought treatment while the rest remained well-irrigated throughout the experiment. Differential recovery by irrigation restoration of the drought treated plants, began on Aug 20’ 2018 (allowing each plant to reach a similar degree of stress) until the experiment end.

### 9. Analyze your data using SPAC Analytics

9.1. Open the SPAC Analytics software. Click on the top right corner to select Control system and the name of the experiment (**Figure 9A**). In the column of the left side of the screen, select “Experiments” (**Figure 9B**) and type the name of the experiment in the Name bar under the Search section. The name of the experiment will appear below the Search section, in the Experiments section (**Figure 9C**). Click on the experiment to open the Info and Plants sections (**Figure 9D**).

**Figure 9.**
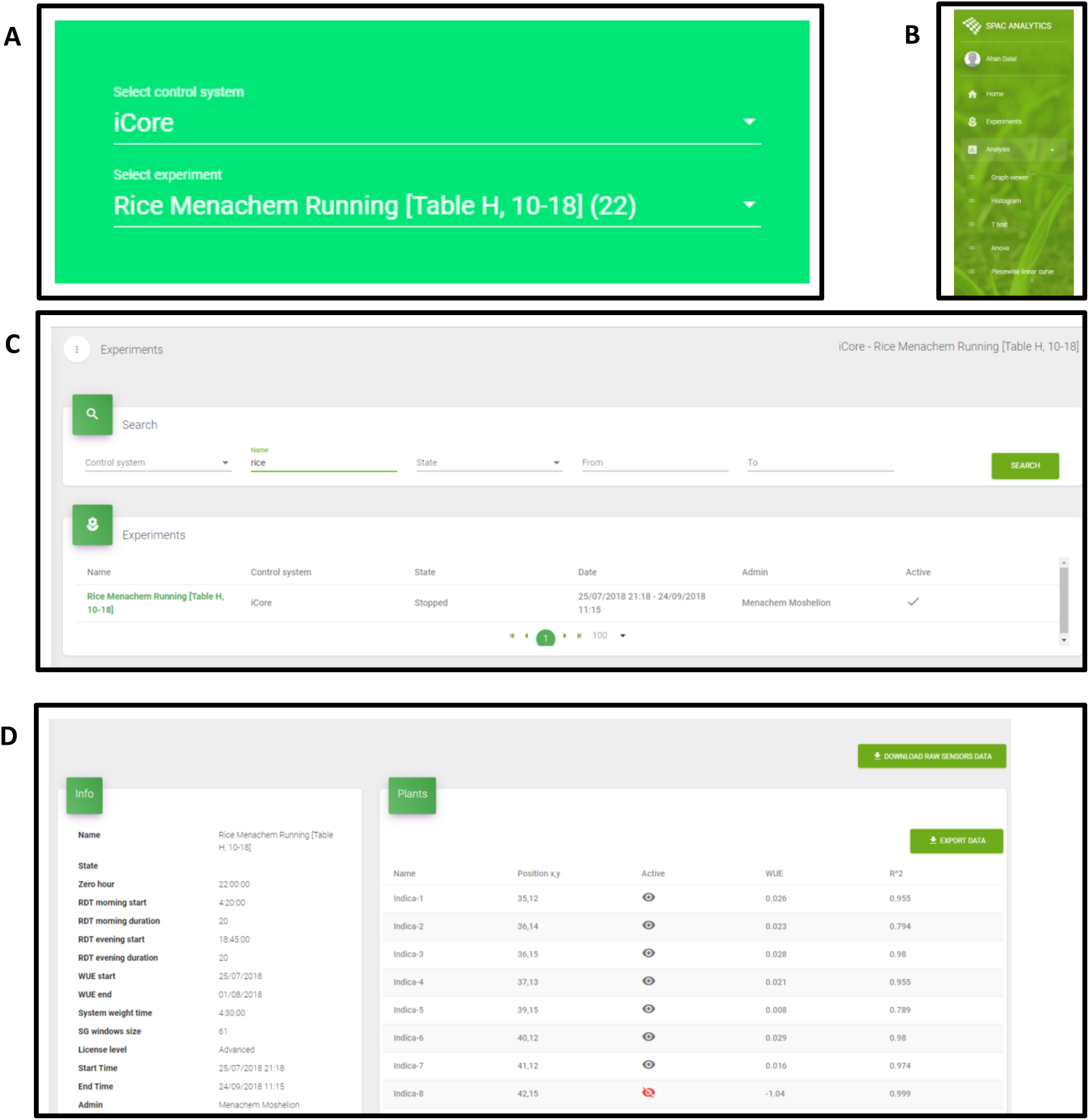
Software windows of “SPAC Analytics” for data analysis.
9.2. In the Info section, edit the WUE start and WUE end dates for a period of at least 3 (preferably more) days before the start of the drought treatment and then click “Update”. The WUE and the *R*^*2*^ value for every pot will appear in the Plants section. You can choose to exclude any scale with a negative WUE value or an *R*^*2*^ value of less than 0.5 by clicking on the eye symbol under the Active column, which will then turn red. This will exclude the selected scale (plant) from all further calculations. The data can be exported by clicking on the Export Data button in the Plants section (**Figure 9D**).
9.3. In the column on the left side of the scree, click on “Analysis”. Different subsections will then appear: Graph viewer, Histogram, T-test, ANOVA and Piecewise linear curve.
9.4. A. Click on “Graph viewer”. In the Filters section, set the dates for your experiment. Click on “Labels” (please see step 7) to select the combination of experimental groups (genotype) and treatment(s). Automatically, all pots in the selected group will appear in the Plant subsection, in which any pots (plants) can be deselected by clicking on them. Up to two different parameters of your choice can be selected at one time as the “Y1 parameter” and “Y2 parameter”. Finally, click on “SHOW GRAPH” (**Figure 8**). B. A line graph of the values of the selected parameter will appear in the Graph Viewer window for each plant. Data from individual plants can be removed from or added to the graph by clicking on their legend symbols on the right of the graph. In the top right corner, there are also options for exporting the data as an Excel file and for enlarging the Graph Viewer window to fill the full screen. More options to modify the graph will appear if you move the cursor to the top right corner of the screen (**Figure 8**).
9.5. The histogram module presents the distribution of a single trait in and between populations for a given time period. To use this module, click on “Histogram”. In the Filters section, set the date and time, parameter, labels and plants as explained in 9.4.A. You can select multiple labels (groups) by clicking on the “+” symbol. Finally, click on “SHOW GRAPH” (**Figure 10**).

**Figure 10.**
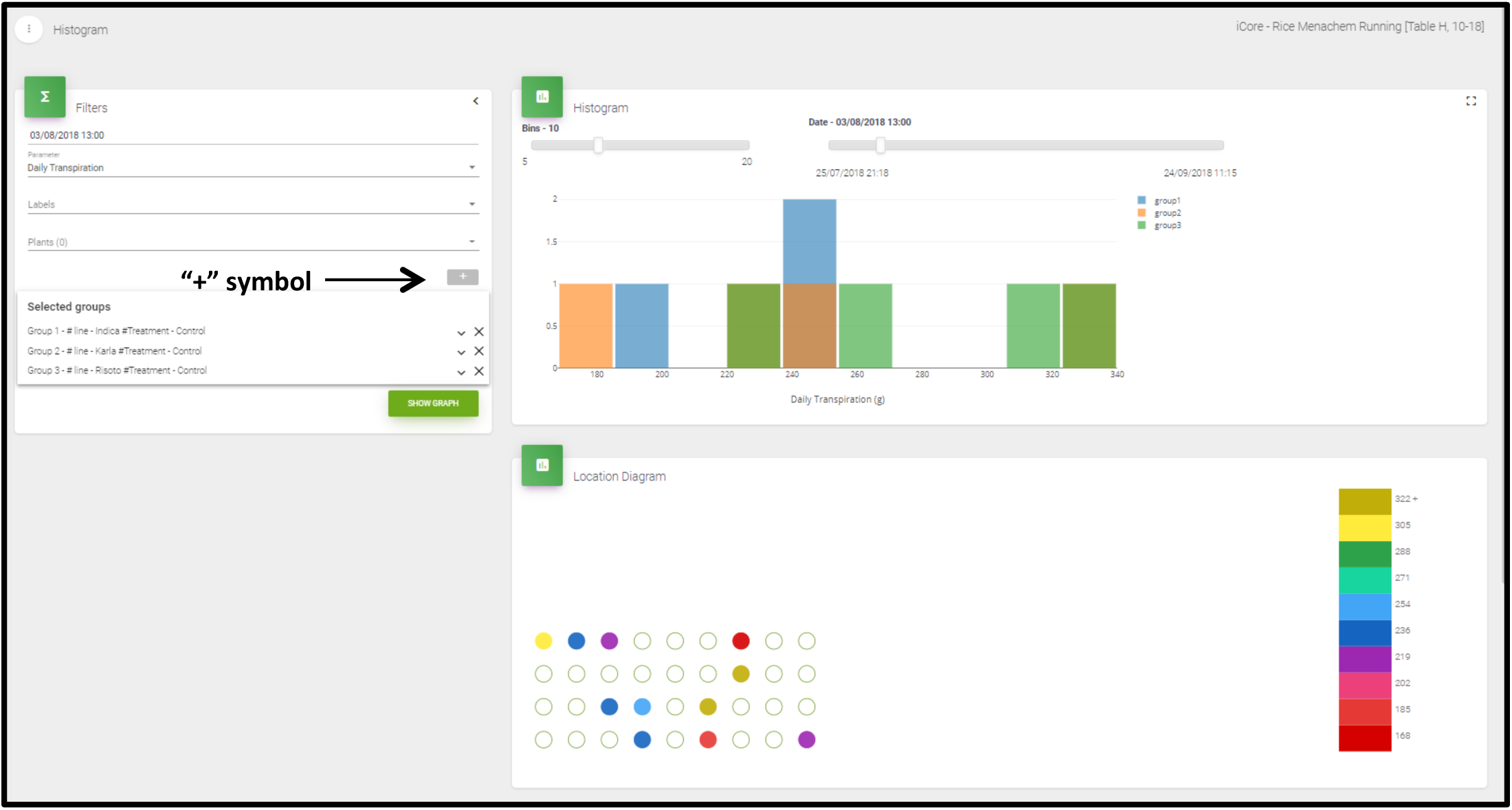
**“**SPAC Analytics” Histogram window. Graphical representation of the of the daily transpiration distribution in the three different rice varieties namely: *Indica, Karla* and *Risotto*, under well irrigated (control) conditions. The histogram will appear in the Histogram section, in which you have the option to change the “Bins” and “Date” at the top of the screen. In the top right corner, there are various options as described in 9.4.B. In the Location Diagram section, the actual location of the plants on the experimental table and their respective trait values can be seen (**Figure 10**).
9.6. Click on “T-test”. To statistically compare the means of any measured trait of two groups, enter the dates, labels, plants and parameters in the T-test Parameters section, as explained in 9.4.A. Set the range of hours to calculate the average values of the data points within your time period of interest. Finally, click on “SHOW GRAPH” (**Figure 11**).

**Figure 11.**
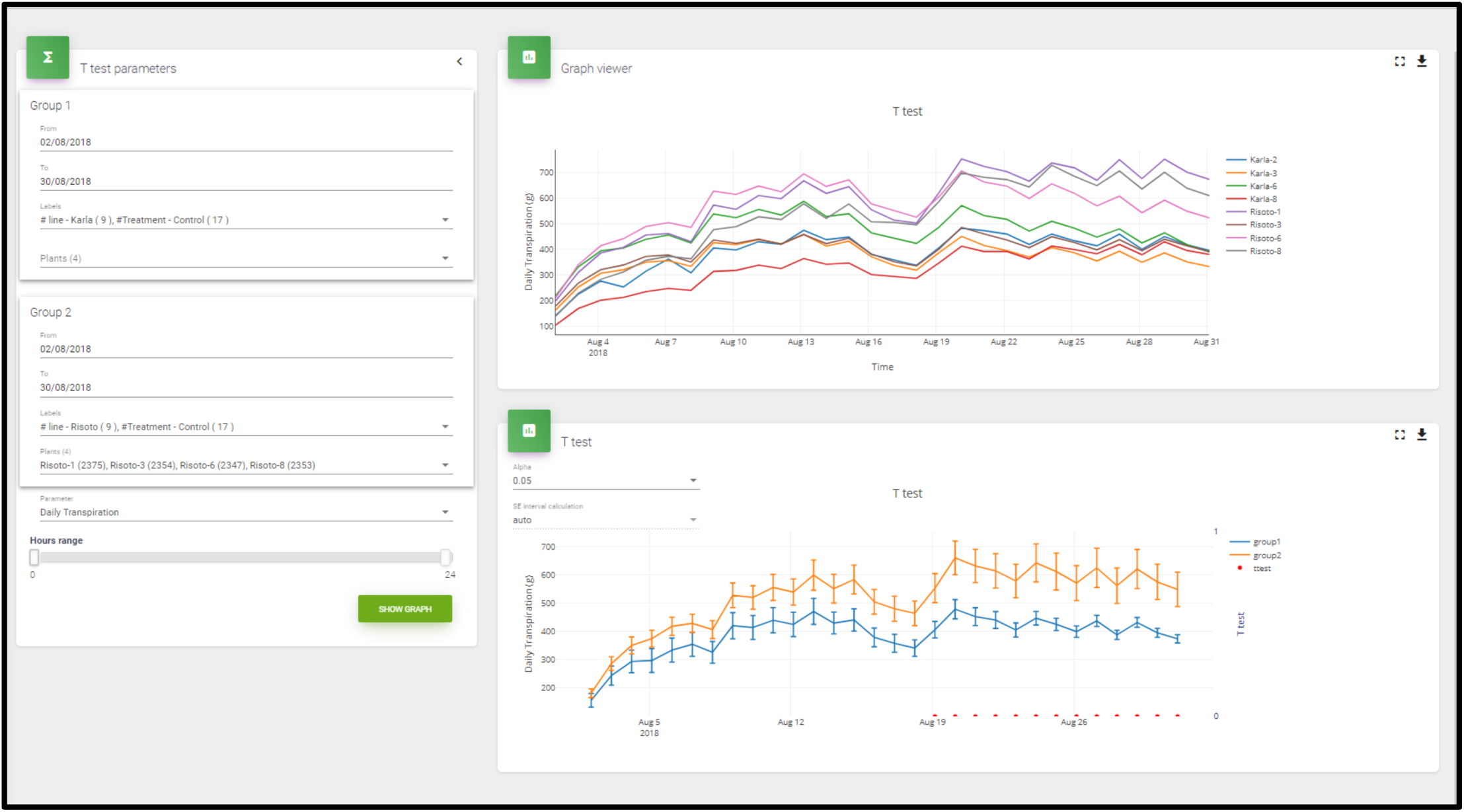
**“**SPAC Analytics” T-test window. Lines represent the difference in daily transpiration, a fundamental and important physiological trait, between two rice varieties namely *Karla* and *Risotto* under well irrigated (control) conditions. The window shows the daily transpiration of the individual plants (top graph) and compares the means ± SE of each group using student’s *t*-test (bottom graph). The statistical analysis is performed automatically by the software. The red dots represent significant differences among treatments (Student’s *t*-test; *p* < 0.05). Two windows will appear on the right side of the screen. The top one is the Graph Viewer section for all of the plants selected from both groups. Below that window is the T-test section, in which will appear the comparison of the two groups as the *t*-test of the physiological parameter selected. Levels of significance can be adjusted by changing the α value in the top left corner of the screen. A red dot will appear under values that are significantly different. In the top right corner, you can view various options, as described in 9.4.B (**Figure 11**).
9.7. Click on “ANOVA”. To statistically compare the means of any measured trait of more than two groups, enter the dates, labels, plants and parameters in the Filters section, as explained in 9.4.A. You can select multiple labels (groups) by clicking on the “+” symbol (as in 9.5). Set the range of hours. Finally, click on “SHOW GRAPH” (**Figure 12**).

**Figure 12.**
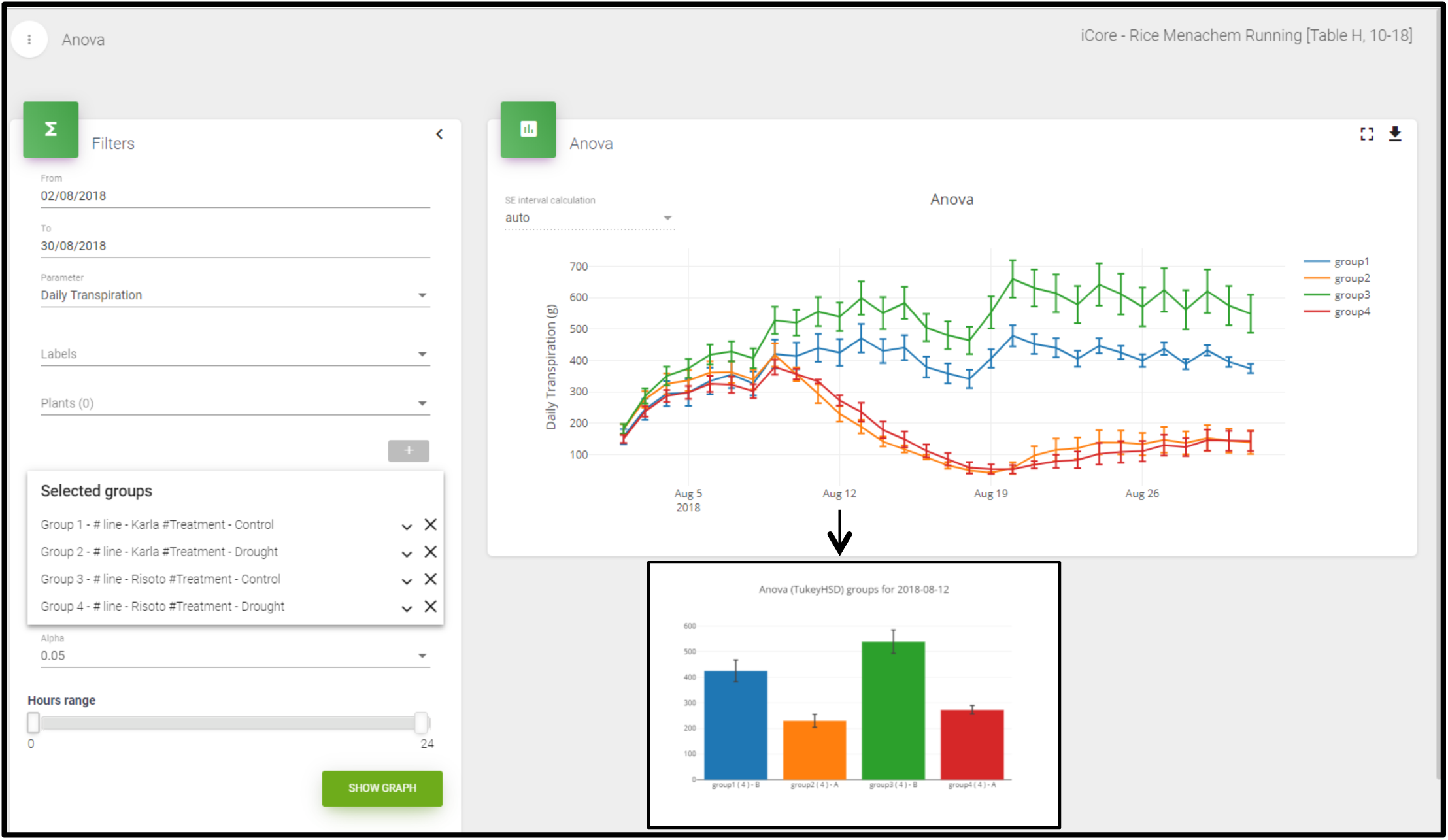
**“**SPAC Analytics” ANOVA window. Graphical representation of the difference in daily transpiration, between two rice varieties *Karla* and *Risotto* under two conditions, well irrigated (control) and drought conditions. The graph presents the daily transpiration as the mean ± SE of each group/treatment compared using ANOVA (Tukey-HSD; *p* < 0.05). The statistical analysis is performed by the software. In the ANOVA section, you can use an ANOVA test (Tukey’s HSD) to compare the physiological parameters of the different groups. Bars represent the standard errors (±SE). In the top right corner of the screen, there are various options as described in 9.4.B. Click on the line graph to view a bar graph comparison for a particular day. Different letters indicate groups that are significantly different from one another (**Figure 12**).
9.8. Presenting the relationship between whole-plant transpiration kinetics or stomatal conductance and VWC is a more accurate way to compare the physiological responses of plants to drought, as compared to a time-based approach. This relationship can be presented using the Piece-wise Linear Curve function.

Click “Piecewise linear curve”. Enter the dates, labels, plants and parameters (both the x-axis and the y-axis) and then set the range of hours in the Filters section, as explained above. The “from” date should be as close as possible to the treatment start date. Set the x-axis parameter to be VWC and the y-axis parameter as the physiological parameter of your choice (e.g., transpiration rate, stomatal conductance, etc.). Finally, click on “SHOW GRAPH”. In the Filter section, click on “Select all recommendations” and then click on “SHOW GRAPH” (**Figure 13**).

**Figure 13.**
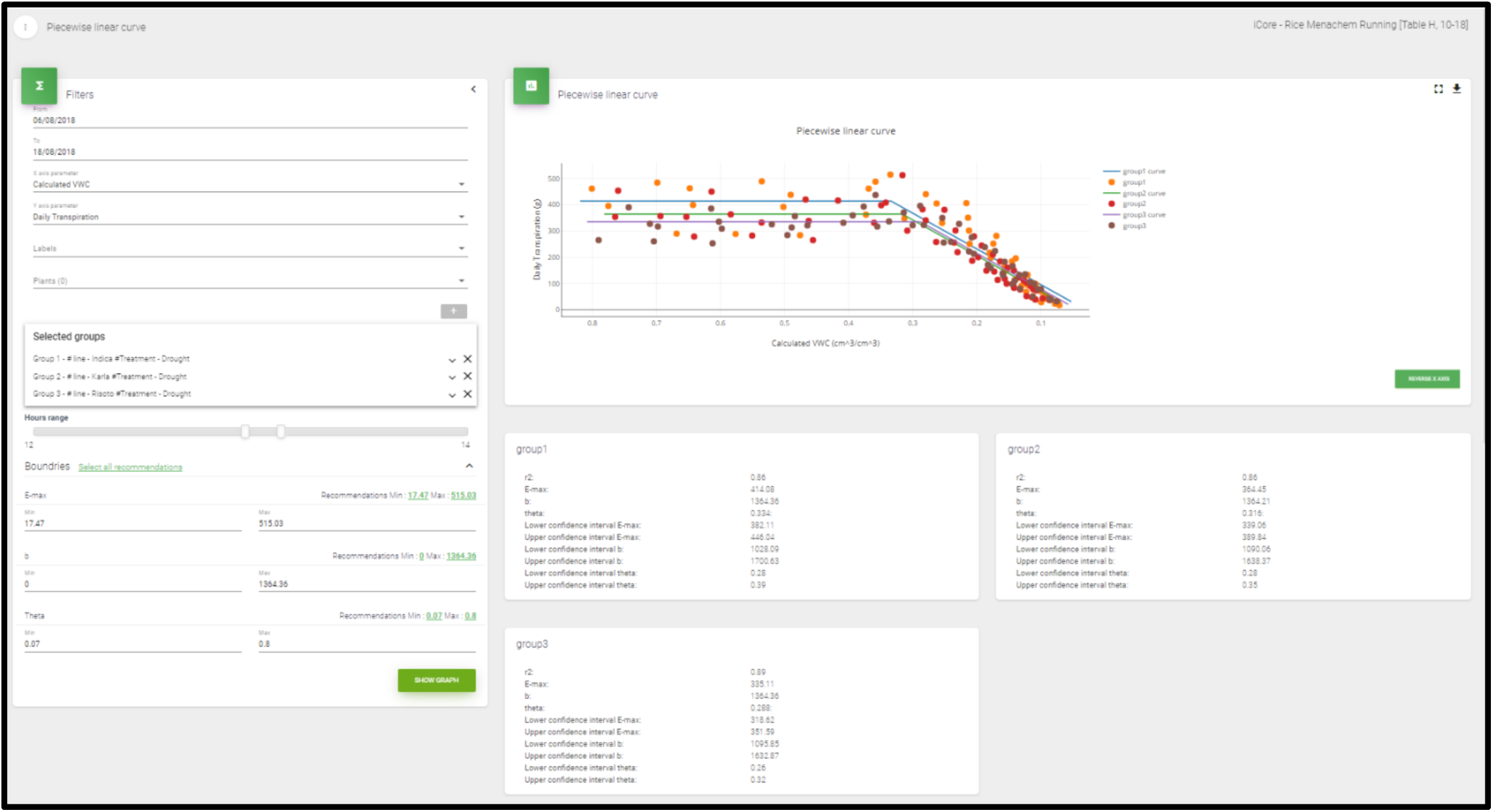
“SPAC Analytics” Piecewise linear curve window. The piecewise linear curve of three rice varieties: *Indica, Karla* and *Risotto* under drought conditions. The software performs the Piecewise linear fit between any physiological parameter (here it is daily transpiration) and the calculated volumetric water content (VWC) for the plants subjected to drought treatment.

## REPRESENTATIVE RESULTS

The duration of our experiment was 29 days. The experiment was conducted in August, when the local weather is warm and stable and the days are long. Two different irrigation scenarios were used to demonstrate the capability of the phenotyping platform for comparing the physiological behavior of three different varieties of rice (i.e., Indica, Karla and Risotto) in the presence of drought stress. There were two drought-stress treatments: (i) optimal irrigation (control) and (ii) a drought that started 5 days after the experiment started, lasted for 14 days, and was followed by a 10-day recovery period (optimal irrigation, Days 19–29). For the sake of simplicity, not all of the varieties and groups are shown in figures. The results show that the high-throughput array system can efficiently measure changes in atmospheric conditions, the soil and the physiology of plants.

### Environmental conditions

Environmental conditions [photosynthetically active radiation (PAR) and vapor pressure deficit (VPD)] were monitored throughout the experiment by an atmospheric probe. The collected data indicated that fluctuations in PAR and VPD during each of the days of the experiment were moderate (**Figure 14**).

**Figure 14.**
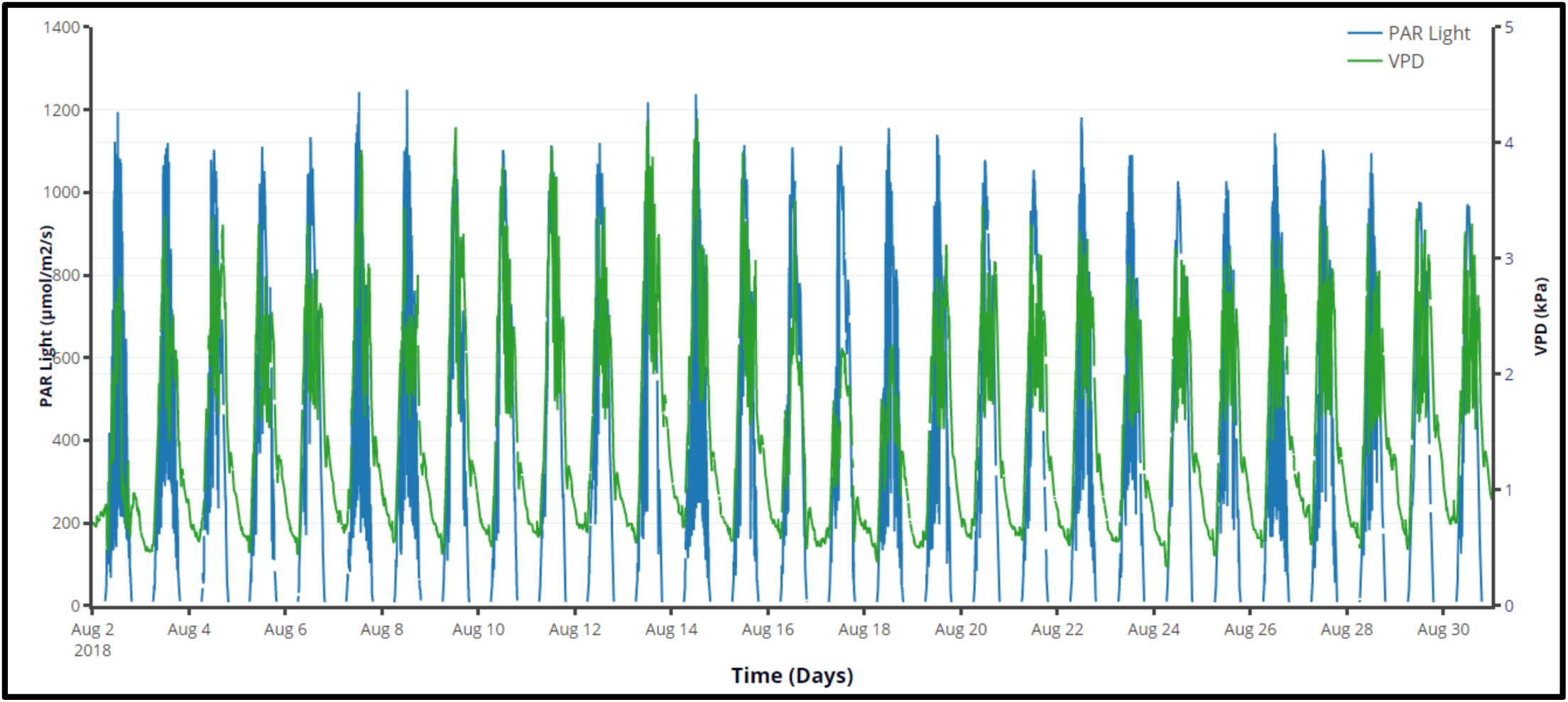
Atmospheric conditions throughout the experiment. The right Y-axis shows the daily vapor pressure deficit (VPD) and the left Y-axis shows the photosynthetically active radiation (PAR) during 29 consecutive days of the experiment. The graph is directly procured from the “SPAC Analytics” software.

The VWC of the drought-treated pots was measured by soil probes throughout the experimental period. The VWC data collected from one drought-treated Indica plant is plotted in **Figure 15**.

**Figure 15.**
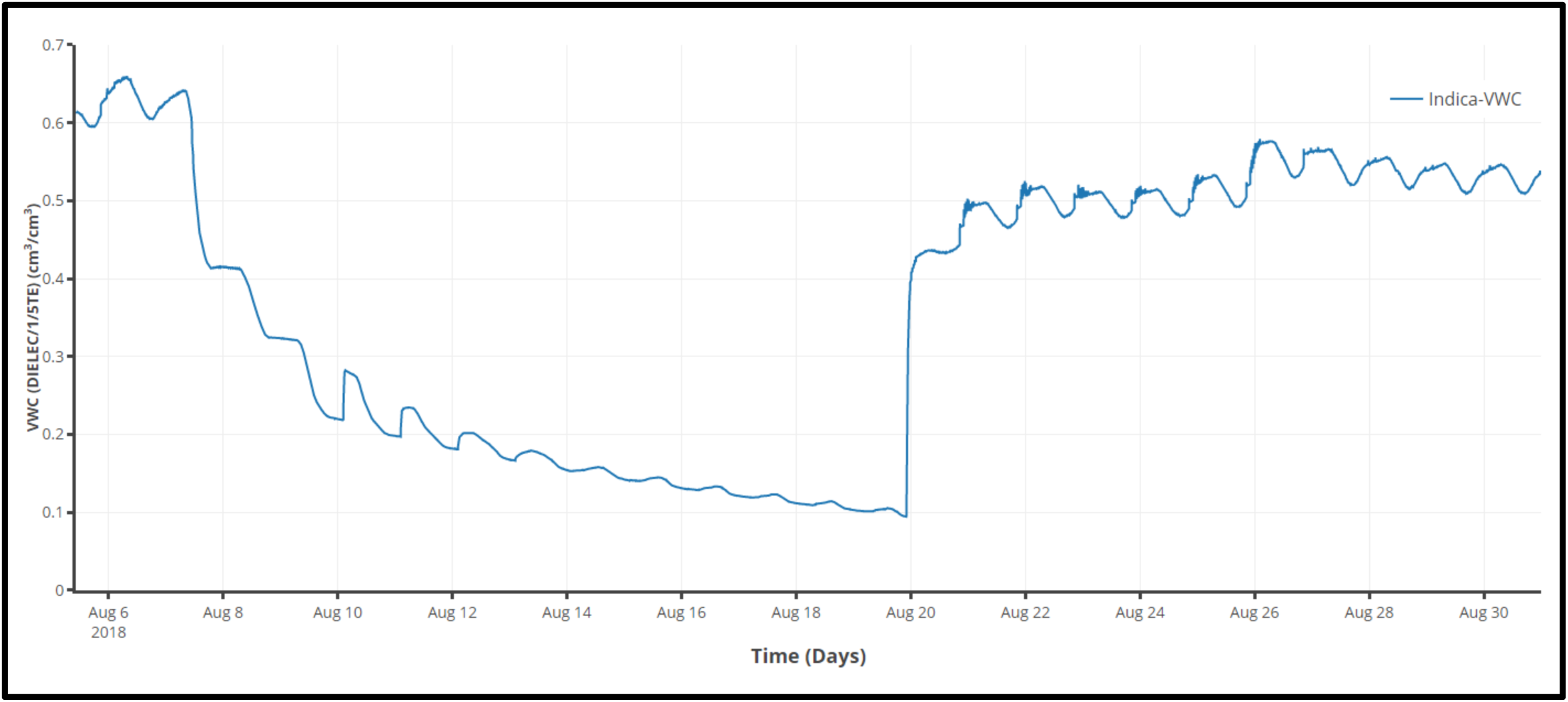
The volumetric water content (VWC) measured by soil probe throughout the experiment. The data represents the VWC of one plant from *Indica* variety that had undergone drought. The graph is directly procured from the “SPAC Analytics” software.

### Physiological parameters

The daily transpiration gradually increased in all four treatments (Karla-control, Karla-drought, Risotto-control and Risotto-drought) during the first stage of the experiment, during which all of the plants were well-irrigated. Later, there was a reduction in transpiration that was associated with the drought period (Day 5 to Day 18) in the two water-deprived treatments. Subsequently, during the recovery period (from Day 18 onward), the daily transpiration increased again in the two water-deprived groups, but to a much lower level than that observed before the drought treatment (**Figure 16**).

**Figure 16.**
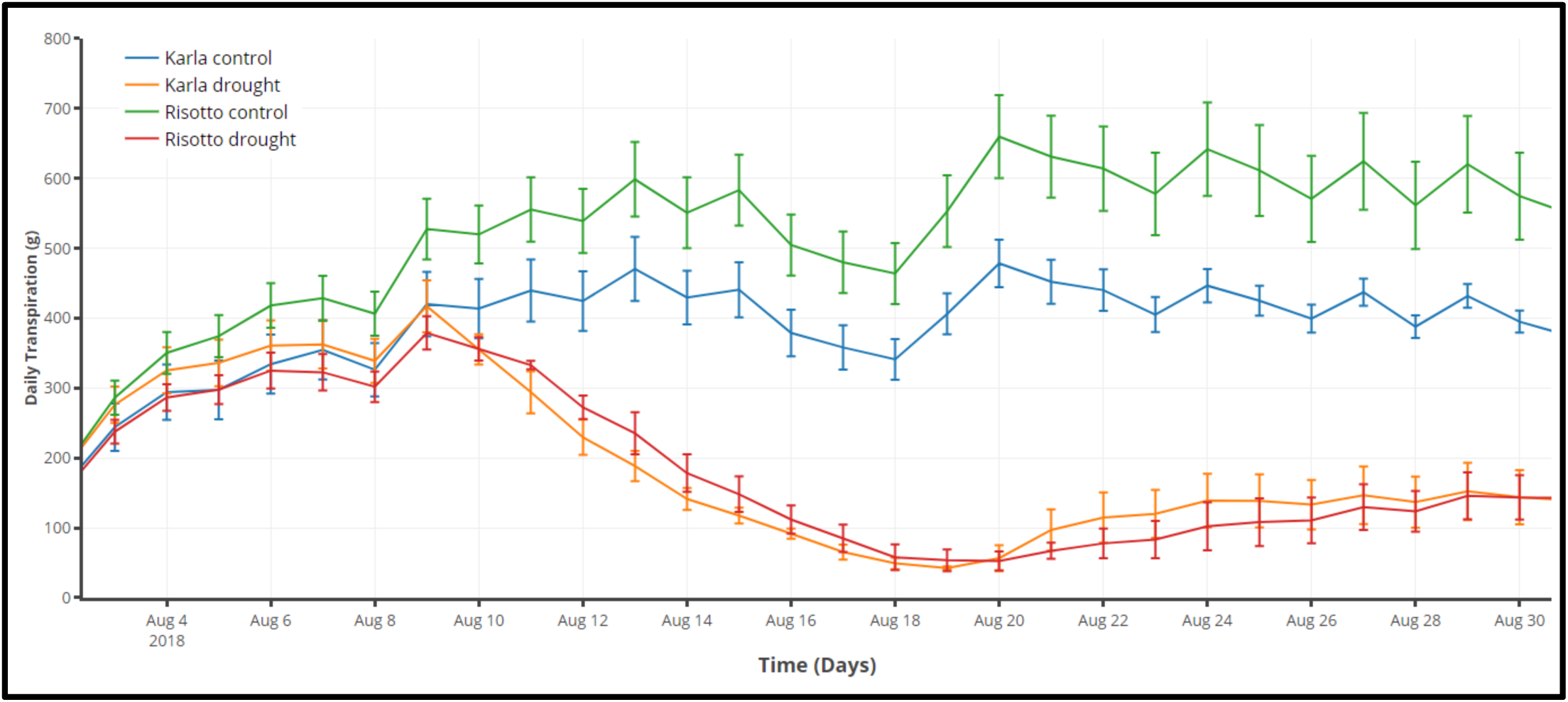
Mean ± SE continuous daily whole-plant transpiration during the entire experimental period. Two varieties, *Karla* and *Risotto* under two treatments (i) well irrigated (control) and (ii) drought. Groups are compared using ANOVA (Tukey-HSD; *p* < 0.05). Each mean ± SE is from at least four plants per group. The graph along with the statistical analysis is directly procured from the “SPAC Analytics” software.

The mean calculated plant weight (i.e., rate of plant weight gain) increased consistently among both the Karla*-*control and the Karla-drought treatments during the first stage of the experiment, when all of the plants received similar irrigation (Days 1–5). When the drought treatment was applied to the Karla plants (Days 5–18), those plants stopped gaining weight and did not resume gaining weight until the recovery stage. At that point, there was an increase in weight that proceeded more slowly than what was observed for the control. In contrast, the weights of the Karla*-*control plants increased continuously throughout the experimental period (**Figure 17**).

**Figure 17.**
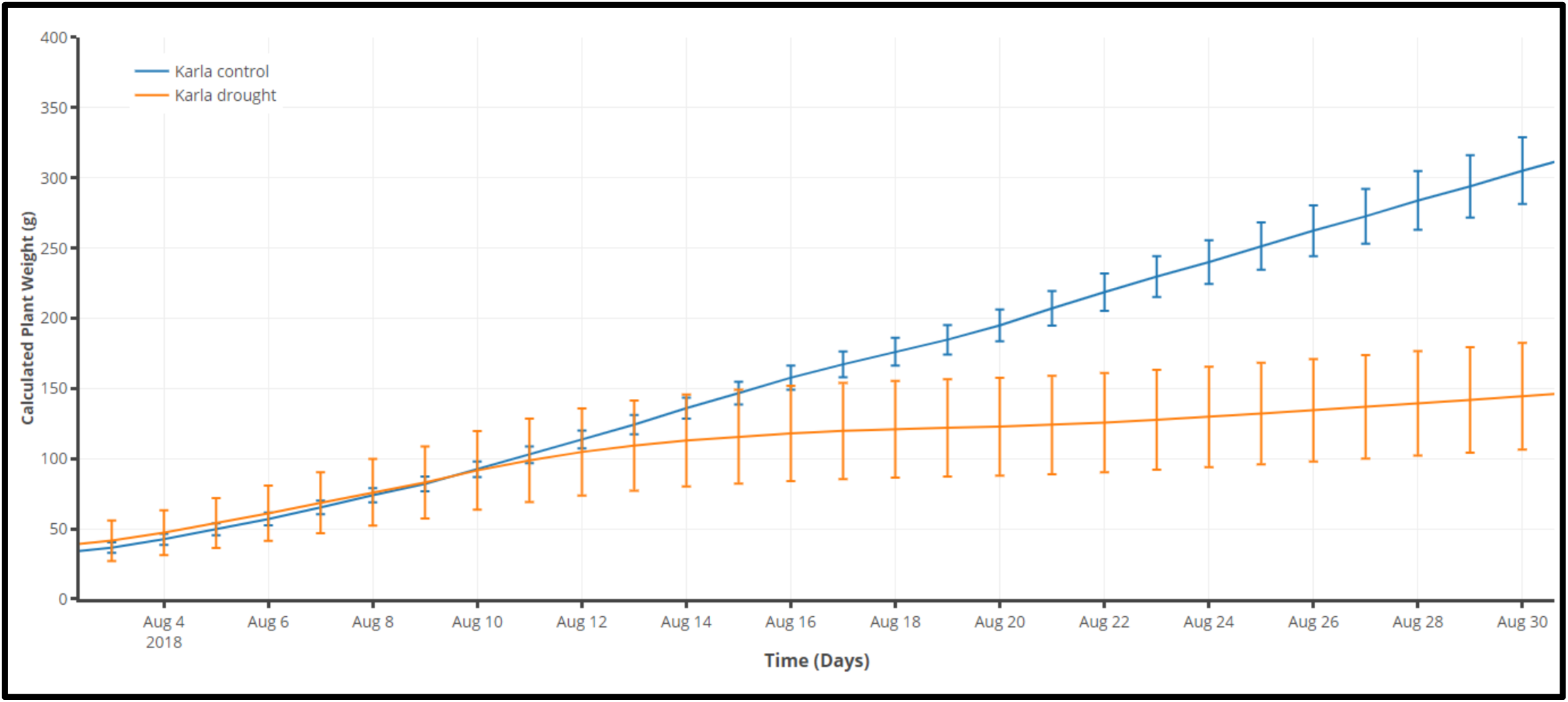
Mean ± SE calculated whole-plant weight during the entire experimental period. The variety *Karla* under (i) well irrigated (control) and (ii) drought scenarios is shown. Groups are compared using ANOVA (Tukey-HSD; *p* < 0.05). Each mean ± SE is from at least four plants per group. The graph along with the statistical analysis is directly procured from the SPAC Analytics software.

**Table 2.** General characteristics of 9 soil types and their compatibility to the gravimetric platform measurements. The measurements were taken using 4L pots, filled with 3.2 L soil media at field capacity (pot capacity). Data are shown as means ± SE. Different letters in the columns indicate significant differences between the medium types, according to Tukey’s HSD test (*P* < 0.05) (*n*>3).

| Soil media type<br>Parameters |  | Coarse sand | Fine sand | Peat based soil | Loamy soil | Vermiculite | Perlite | Compost | Porous Ceramic Small - sized | Porous Ceramic Mix-sized |
| --- | --- | --- | --- | --- | --- | --- | --- | --- | --- | --- |
| Total available water (TAW, ml) | | 860 ± 7.2 (F) | 883.1 ± 24 (F) | 1744.3 ± 8.2 (B) | 1076.3 ± 35.9 (E) | 1286 ± 22.4 (D) | 1119.9 ± 8.5 (E) | 2089.6 ± 61.6 (A) | 1713 ± 25.9 (B) | 1503.6 ± 15.4 (C) |
| Volumetric water content (VWC, ml<sup>3</sup>/ml<sup>3</sup>) |  | 0.26 (F) | 0.27 (F) | 0.54 (B) | 0.33 (E) | 0.4 (D) | 0.35 (E) | 0.65 (A) | 0.53 (B) | 0.46 (C) |
| Bulk density (BD, g/cm<sup>3</sup>) |  | 1.7 (A) | 1.6 (B) | 0.2 (G) | 1.5(C) | 0.2 (F) | 0.1 (H) | 0.1 (G) | 0.7 (E) | 0.8 (D) |
| Soil weight stability (SWS, g/d) | | ±2.3 ± 0.3 (B) | ±4.3 ± 0.3 (B) | ±6.7 ± 0.8 (B) | ±2.9 ± 0.9 (B) | ±7.6 ± 2.8 (B) | ±14.9 ± 0.7 (A) | ±4.3 ± 1.2 (B) | ±1.9 ± 0.4 (B) | ±1.3 ± 0.1 (B) |
| Soil weight stability with reserved water (please see section 6.14) in the bath (g/d) | | 3 ± 0.4 (B) | 3.3 ± 0.4 (B) | 10.6 ± 3 (A) | 3.2 ± 1.2 (B) | 2.7 ± 0.8 (B) | 6.3 ± 0.5 (A) | 1.5 ± 0.3 (B) | 1.9 ± 0.3 (B) | 1.6 ± 0.3 (B) |
| Pot capacity gravimetric moisture content (SWC, please see section 8.2 ) |  | 0.18 (G) | 0.23 (G) | 4.25 (B) | 0.23 (G) | 3.0 (D) | 3.79 (C) | 6.13 (A) | 0.99 (E) | 0.74 (F) |
| Relative drainage capability |  | Excellent | Medium | Low | Medium-low | Excellent | Excellent | Medium | Excellent | Excellent |
| Relative time to reach pot capacity |  | Fast | Fast | Slow | Fast | Slow | Slow | Slow | Fast | Fast |
| Relative cation exchange capacity (CEC) |  | Low | Low | High | Low | High | Low | High | High | High |
| Compatibility to: | Root washing (at the end of the experiment) | ++ | ++ | - | + | + | ++ | - | ++ | ++ |
|  | Nutrient/biostimulant treatment | ++ | ++ | - | - | + | ++ | - | ++ | ++ |
|  | Salinity experiment | ++ | ++ | + | + | + | ++ | - | ++ | ++ |
|  | Growth rates measurement accuracy | +++ | ++ | + | + | + | -,+ | + | +++ | ++ |
|  | Physical soil structure recovery after drought | +++ | +++ | -,+ | ++ | - | + | - | +++ | +++ |

\* Total available water (TAW) = soil wet weight (at pot capacity) – soil dry weight; Volumetric water content (VWC) = TAW/soil volume; Bulk density (BD) = soil dry weight/soil volume; Soil weight stability (SWS) = Average of delta between RDT morning of 4 consecutive days (media at pot capacity with no-plant); Pot capacity gravimetric moisture content (SWC), for calculation please see section 8.2.

## DISCUSSION

The genotype–phenotype knowledge gap reflects the complexity of genotype × environment interactions (reviewed by Dalal et al 2017; Gosa et al 2018). It might be possible to bridge this gap through the use of high-resolution, high-throughput diagnostic and phenotypic screening platforms to study whole-plant physiological performance and water-relation kinetics (Moshelion and Altman, 2015; Dalal et al 2019). The complexity of genotype × environment interactions makes phenotyping a challenge, particularly in light of how rapidly plants respond to their changing environments. Although various phenotyping systems are currently available, most of them are based on remote sensing and advanced imaging techniques. Although those systems provide simultaneous measurements, to a certain extent, their measurements are limited to morphological and indirect physiological traits (Araus et al., 2018). Physiological traits are very important in the context of responsiveness or sensitivity to environmental conditions (Ghanem et al., 2015). Therefore, direct measurements taken continuously and simultaneously at a very high resolution (3-min intervals) can provide a very accurate description of a plant’s physiological behavior.

Any phenotyping system designed to address plant–environment interactions should provide tools to standardize the applied stresses, create a truly randomized experimental structure, minimize pot effects and compare multiple dynamic behaviors of plants under changing environmental conditions. The high-throughput functional phenotyping approach described in this paper addresses those issues as noted below.

1. In order to correlate the plant’s dynamic response with its dynamic environment and capture a complete, big picture of complex plant–environment interactions, both environmental conditions (**Figure 14**) and plant responses (**Figure 16**) must be measured continuously. This method enables the measurement of physical changes in the growing medium and atmosphere continuously and simultaneously, alongside plant traits (soil–plant–atmosphere-continuum, SPAC).
2. To best predict how plants will behave in the field, it is important to perform the phenotyping process under conditions that are as similar as possible to those found in the field (Gosa et al., 2018). We conduct our experiments in a greenhouse under semi-controlled conditions to mimic field conditions as much as possible.
3. Field phenotyping and greenhouse phenotyping (pre-field) have their own objectives with different experimental set-ups. Pre-field phenotyping assists the selection of promising candidate genotypes that have a higher probability of doing well in the field, to help make field trials more focused and cost-effective. However, pre-field phenotyping involves a number of limitations (e.g., pot effects) that can cause plants to perform differently than they would under field conditions (Sinclair et al., 2017; Gosa et al., 2019). Small pot size, water loss by evaporation and heating of the lysimeter scales are examples of factors in greenhouse experiments that may lead to pot effects (reviewed by Gosa et al., 2019). The method described here is designed to minimize these potential effects in the following manner:

a. The pot size is chosen based on the genotype.
b. The pots and lysimeter scales are insulated to prevent heat from being transferred and any warming of the pots.
c. This system involves a carefully designed irrigation and drainage system.
d. There is a separate controller for each pot, to enable true randomization with self-irrigating and self-monitored treatments
4. This system involves direct physiological measurements at field-like plant densities, which eliminates the need for either large spaces between the plants or moving the plants for image-based phenotyping.
5. This system includes real-time data analysis, as well as the ability to accurately detect the physiological stress point (θ) of each plant. This enables the researcher to monitor the plants and make decisions, in execution and/or sample collection, while the experiment is ongoing.
6. The system’s easy and simple weight calibration facilitates efficient calibration.
7. High-throughput systems generate massive amounts of data, which present additional data-handling and analytical challenges (Houle et al., 2010; Fiorani and Schurr, 2013). The real-time analysis of the big data that is directly fed to the software from the controller is an important step in the translation of data into knowledge (Negin and Moshelion, 2017) that has great value for practical decision-making.

## CONCLUSION

This high-throughput physiological phenotyping method might be helpful for conducting greenhouse experiments with near-to-field conditions. The system is able to measure and directly calculate water-related physiological responses of plants to their dynamic environment, while efficiently overcoming most of the problems associated with pot experiments. This system’s abilities are extremely important in the pre-field phenotyping stage, as they offer the possibility to predict yield penalties during the early stages of plant growth.

## ACKNOWLEDGMENTS

This work was supported by the ISF-NSFC joint research program (grant No. 2436/18) and was also partially supported by the Israel Ministry of Agriculture and Rural Development (Eugene Kandel Knowledge Centers) as part of the Root of the Matter – The Root Zone Knowledge Center for Leveraging Modern Agriculture.

## DISCLOSURES

The authors have nothing to disclose.

